# Genome-wide chromatin accessibility analyses provide a map for enhancing optic nerve regeneration

**DOI:** 10.1101/2021.02.25.432980

**Authors:** Wolfgang Pita-Thomas, Tassia Mangetti Gonçalves, Guoyan Zhao, Valeria Cavalli

## Abstract

Retinal Ganglion Cells (RGCs) lose their ability to grow axons during development. Adult RGCs thus fail to regenerate their axons after injury, leading to vision loss. To uncover mechanisms that promote RGC axon regeneration, we identified transcription factors (TF) and chromatin accessible sites enriched in embryonic RGCs (high axon growth capacity) compared to postnatal RGC (low axon growth capacity). Developmental stage-specific gene expression changes correlated with changes in promoter chromatin accessibility. Binding motifs for TFs such as CREB, CTCF, JUN and YY1 were enriched in the differentially opened regions of the chromatin in embryonic RGCs and proteomic analysis confirmed their expression in RGC nuclei. The CREB/ATF binding motif was widespread at the open chromatin region of known pro-regenerative TFs, supporting a role of CREB in regulating axon growth. Consistently, overexpression of CREB fused to the VP64 transactivation domain in RGCs promoted axon regeneration after optic nerve injury. Our study provides a map of the chromatin accessibility during RGC development and highlights that TF associated with developmental axon growth can stimulate axon regeneration in mature RGC.

## INTRODUCTION

Retinal Ganglion Cells (RGCs) in the retina receive visual information from bipolar cells and transmit it to several regions in the brain via axons projecting into the optic nerve. RGCs originate from retinal precursor cells and most of them are born between E11 and E20 (Rapaport *et al*, 2004). Transcription factors (TFs) such as Pax6, Math5, Pou4f2, form a hierarchical network that regulates RGC differentiation from retinal precursors in space and time. Once differentiated, RGCs grow their axons towards the brain through the optic nerve. Early born RGCs (born before E16) arrived at the superior colliculus before birth while late born RGCs (born after E16) arrive at the superior colliculus at P4/P5 (Dallimore *et al*, 2002). Right before eye-opening (P12), all RGCs have formed stable synapses with their brain targets, providing a functional visual system. The maturation of the visual system correlates with a decrease in the regenerative capacity of RGCs. Therefore, RGC injury in adult mammals is followed by axon regeneration failure and a degeneration process that leads to cell death. As a result, diseases that cause optic nerve damage such as traumatic optic neuropathy, glaucoma, or optic nerve ischemia result in an irreversible loss of vision. Identifying molecular and cellular mechanisms that increase survival and regeneration of RGC may offer new treatment strategies for patients with glaucoma or other types of optic neuropathies.

RGCs lose their capacity to grow axons after birth (Goldberg *et al*, 2002b; Steketee *et al*, 2014). Axon growth capacity in rat RGCs peak at embryonic day 21 (E21), and start declining after birth reaching their minimum growth potential at postnatal day 11 (P11), right before eye-opening (Goldberg *et al*., 2002b). These experiments suggested that the axon growth program is substituted by a synapse function program. Multiple transcription factors that promote axon growth, such as SOX11 (Norsworthy *et al*, 2017) and KLF7 (Moore *et al*, 2009) are downregulated during development in RGCs. Other transcription factors that inhibit axon growth such as KLF4 and KLF9 are upregulated during development (Moore *et al*., 2009). Manipulation of these transcription factors promotes axon regeneration after optic nerve injury (Apara *et al*, 2017; Blackmore *et al*, 2012; Lu *et al*, 2020; Moore *et al*., 2009; Norsworthy *et al*., 2017).

Chromatin accessibility may represent another epigenetic mechanism controlling the axon growth program. It has been suggested that chromatin accessibility of some TFs such as KLF7 is limited in mature neurons, and transactivation domains are needed to boost regeneration (Blackmore *et al*., 2012; Venkatesh *et al*, 2018). Proteins that modify chromatin accessibility, such as histone deacetylases and histone acetyltransferase have also been shown to play a role in sensory neuron regeneration (Cho *et al*, 2013; Finelli *et al*, 2013; Hervera *et al*, 2019; Puttagunta *et al*, 2014) and RGCs (Gaub *et al*, 2011; Pita-Thomas *et al*, 2019; Schmitt *et al*, 2014; Schmitt *et al*, 2018). The interplay of TFs and chromatin accessibility regulates the expression levels of downstream genes such as *Gap43* and *Tubb3* whose final location is at the growth cone (Patodia & Raivich, 2012). Some of these downstream genes may also regulate signaling pathways that control metabolism. Activating pathways that control cell growth, endosome recycling, and ribosome biogenesis such as overexpressing protrudin-1 or activating the mTOR pathway promotes abundant axon regeneration (Park *et al*, 2008; Petrova *et al*, 2020; Pita-Thomas *et al*., 2019). Therefore, the complex interaction of genes that control growth cone dynamics and cell metabolism increase the axon growth capacity of RGCs. Altogether, these studies point to the complex network of regenerative associated genes epigenetics, TFs and downstream genes regulating the decline in axon growth capacity in RGCs during development.

Analysis of chromatin accessibility has previously resulted in the prediction of target genes of TFs, the hierarchy of TFs in a given network, and the discovery of TFs with unknown roles in developmental biology (Xie *et al*, 2020). Understanding the hierarchical order of these TFs and unraveling the chromatin accessibility at different developmental stages may allow the reprogramming of adult RGCs for axon growth. Here we characterized the chromatin accessibility changes during the developmental period of axon growth decline in rat RGCs (E21 vs P11) using ATAC-seq (Assay for Transposase-Accessible Chromatin using sequencing) and found that changes at the promoter region correlate with gene expression changes determined using RNAseq. By using a proteomics approach, we also determined the transcription factors expressed in RGC nuclei. Transcription factor binding site (TFBS) enrichment analysis identified CREB as one of the most represented TFs in the promoter regions that change in their accessibility between E21 and P11. Finally, overexpressing CREB fused to a VP64 transactivation domain in RGCs induced axon regeneration after optic nerve injury, mimicking the axon growth capacity of embryonic neurons. Our data provides a map of the chromatin accessibility during RGC development and highlights that manipulating TF associated with developmental stages can stimulate axon growth in adulthood.

## RESULTS

### Genetic programs that control axon growth are downregulated during RGC development

To determine the transcriptional changes that occur between embryonic 21 days (E21) and postnatal 11 days (P11) RGCs, RNA was extracted from immunopanned RGCs and sequenced. RNA expression analysis showed that RGC markers such as *Pou4f1* and *RBPMS* were highly enriched compared to markers of microglial, glial, photoreceptor, bipolar, and amacrine cells (Supplementary Figure 1A) in accordance with previous publications reporting 99% purity of RGCs using this technique (Goldberg *et al*., 2002b). Differential expression analysis (Supplementary File 1) showed that mRNA changes in E21 and P11 samples clustered with their particular developmental stage as expected (Fig 1A and Supplementary Figure 1B). Genes differentially upregulated in E21 (cut off of log_2_ fold change >1 and p adj <0.01) included known pro-regenerative TFs and other RAGs such as *Sox11, Smad1, Klf6, Klf7, Myc, Tubb3,* and *Gap43*, while inhibitory RAGs such as *Klf9* and *Klf2* were downregulated (Fig 1B). GO analysis revealed that Biological Processes (BP) such as axon guidance and axonogenesis are enriched at E21, whereas neurotransmitter transport and regulation of membrane potential are enriched at P11 (Fig 1C). Molecular function at P11 also showed enrichment in ion channels, suggesting a role in synapse function and neuronal activity (Fig 1D). Cellular components (CC) enriched at E21 are predominantly related to “ribosome biogenesis” suggesting active cell growth at this stage while CC at P11 pointed to ion channels and synaptic membrane (Fig 1E). KEGG analysis corroborated that pathways related to ribosome function and axon guidance are enriched in E21 while neurophysiological pathways typical of synapse function are enriched in P11 (Fig 1F). Altogether, this transcriptional analysis reveals that, as expected, E21 RGCs are in an active axon growth stage while P11 RGCs have already established synapses and downregulated the axon growth program.

**Figure 1.**
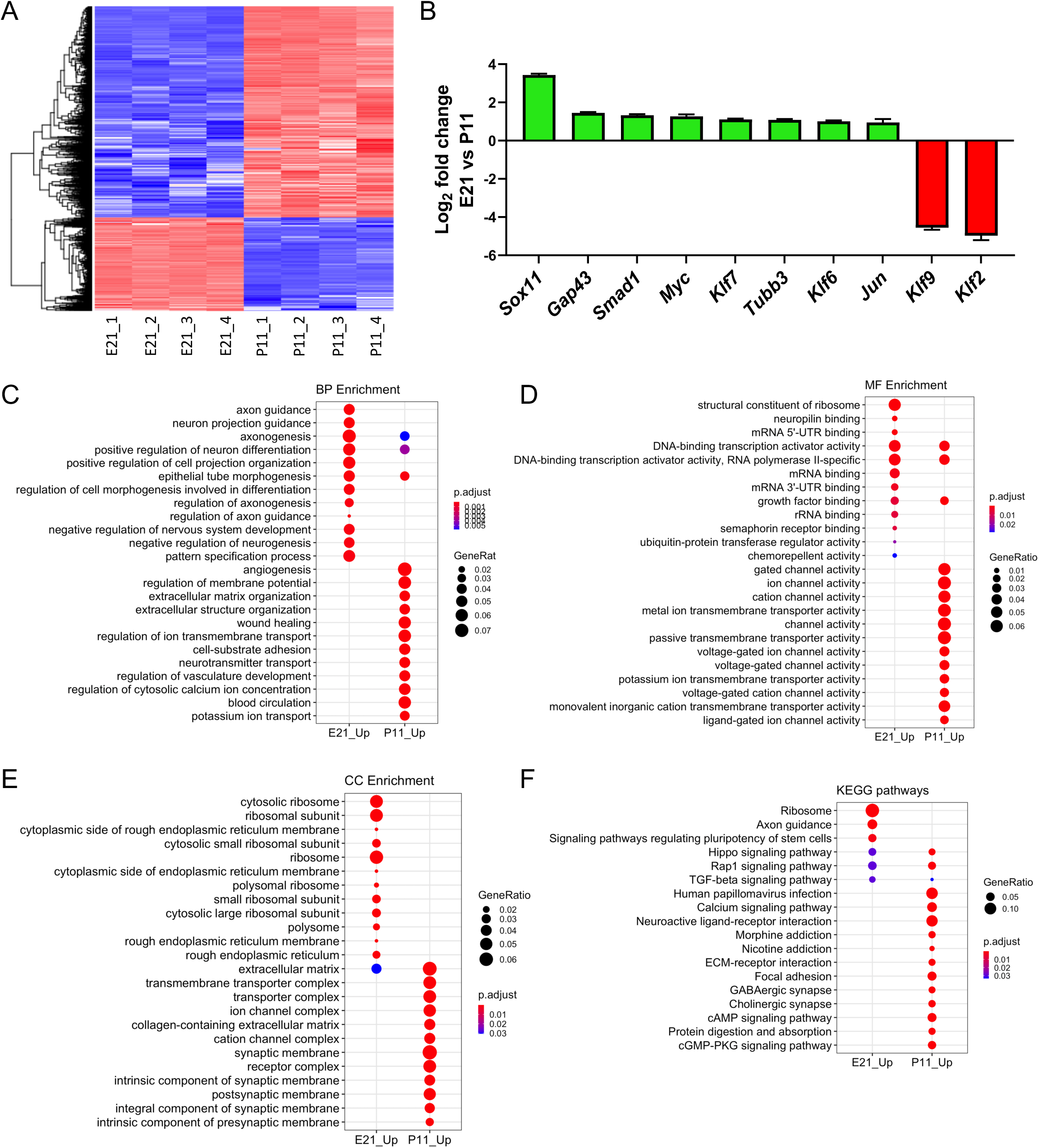
RNAseq analysis of E21 and P11 purified RGCs. A) Genes differentially expressed between E21 and P11 (cut off of log2 fold>1 change and false discovery rate (FDR) adjusted p values < 0.01, which includes a Benjamini-Hochberg correction). n=4 RGC preparations from independent rat litters for each developmental stage. B) Differentially expressed regenerative associated genes described in literature changed consistently with the decline of axon growth capacity between E21 and P11. C) Gene Ontology (GO) analysis of the genes significantly changing between E21 and P11. Twelve most significantly enriched Biological Processes (BP) are represented. D) GO analysis. Twelve most significantly enriched Molecular Functions. E) GO analysis. Twelve most significantly enriched Cellular Components (CC) are represented. F) KEGG analysis of genes significantly changing between E21 and P11. Twelve most significantly enriched pathways are represented. For pathways analysis, FDR corrected P value < 0.05 were considered as significant.

### Changes in chromatin accessibility at the promoter region strongly correlate with changes in gene expression during RGC development

To determine the changes in chromatin accessibility between embryonic and postnatal RGCs, immunopurified rat E21 and P11 RGCs were processed for ATAC-seq (n=2). DNA libraries were sequenced generating 40.9 ± 3.6 million reads. ATAC-seq Integrative Analysis Package (AIAP) was used for sequence quality assurance, mapping, peak calling, and downstream differential open chromatin analysis (Liu *et al*, 2020). We identified 150 ± 17.3 thousand open chromatin regions (OCRs) that were called by MACS2 for each sample (Zhang *et al*, 2008). PCA analysis of the OCRs present in each of the four samples was performed showing that embryonic and postnatal replicates clustered separately (Fig S2A). Pearson correlation coefficient (PCC) between the per-genomic region read count vectors at 1kb and 5kb resolution for each pair of samples was used to assess the global similarity between biological replicates. Each developmental stage were highly correlated between replicates (R>0.96, Supplementary Figure 2B) and were enriched near transcription starting sites (TSS) (Figure 2B), highlighting the quality and reproducibility of the data. The distance to the nearest TSS and distribution of all the OCRs in genomic features (promoters, UTR, exon, intron, downstream, and intergenic) were similar among all the samples (Supplementary Figure 2C).

**Figure 2.**
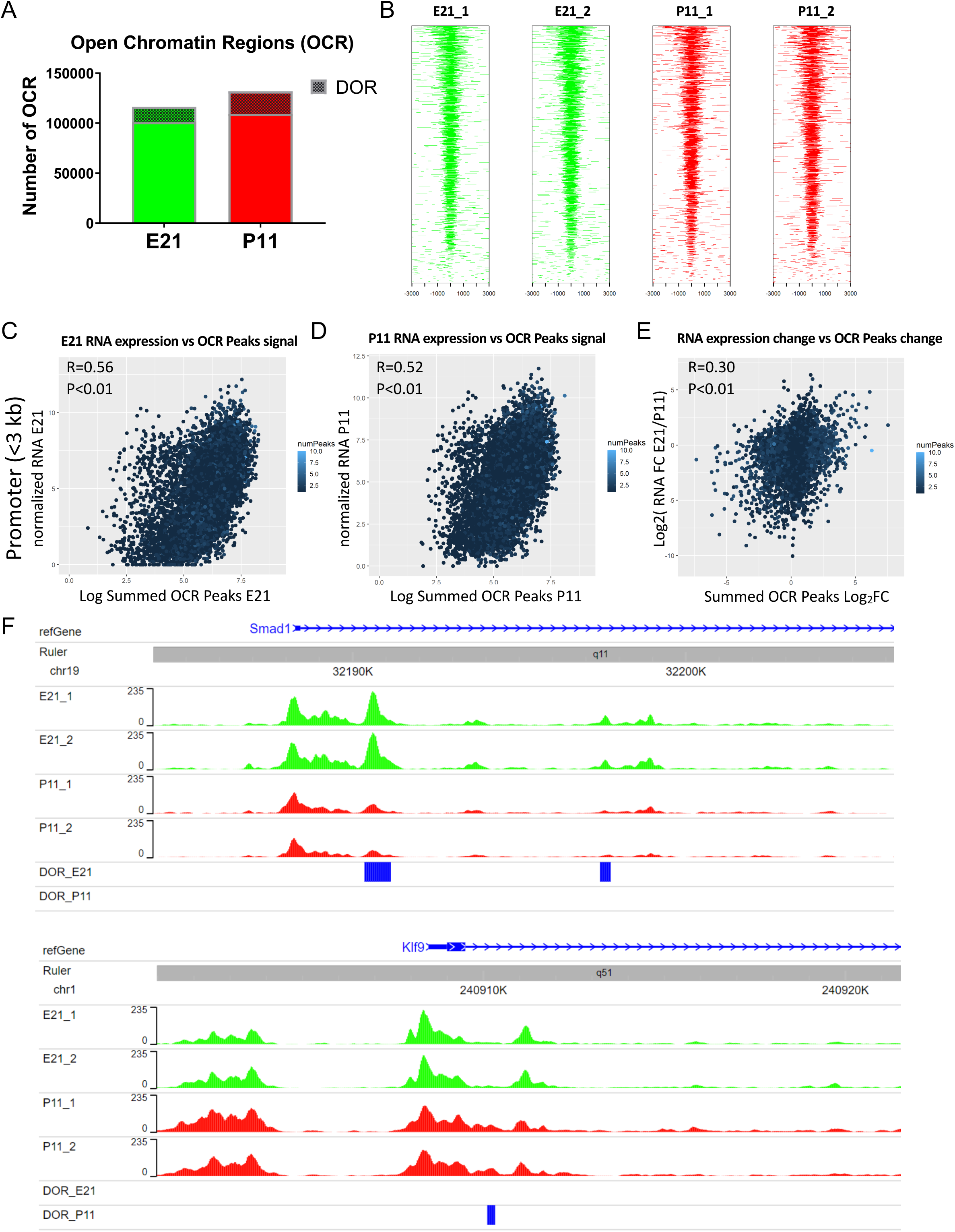
Chromatin accessibility changes at the promoter region (<3 kb from Transcription Start Site) correlate with RNA gene expression changes. A) Number of open chromatin regions (OCRs) at E21 and P11. N=2 RGC samples from independent rat litters for each developmental stage. Cross-hatched area corresponds to the number of differentially opened regions (DORs). DORs were obtained using DESeq2 (Love *et al*., 2014) implemented in the ATAC-seq Integrative Analysis Package (AIAP) package with default parameters. B) Distribution of OCRs with respect to the closest gene Transcription Starting Site (TSS) in the 4 samples. C) Pearson correlation coefficient (p<0.01) between E21 gene RNA expression and chromatin accessibility (sum of all peak values) at the promoter region of genes in E21. D) Pearson correlation coefficient (p<0.01) between P11 gene RNA expression and chromatin accessibility at the promoter region of genes in P11. E) Pearson correlation coefficient (p<0.01) between the change in RNA expression between E21 and P11 and the change in chromatin accessibility at the promoter region of genes between those stages. F) Visual representation of chromatin accessibility of E21 and P11 replicates near the TSS of regenerative associated genes *Smad1* (promoter of axon growth) and *Klf9* (inhibitor of axon growth). DORs are represented in blue accordingly to the developmental stage where this region is more accessible.

A total of 116,417 OCRs were shared by both E21 samples, and 131,806 OCRs in both P11 samples (Fig 2A). We merged all the peaks to generate the final peaks from all samples and determined whether mRNA expression levels correlate with chromatin accessibility at promoter region (<3 kb from TSS), genic region (intron and exon), and distal region (>3 kb but <100 kb from TSS) at each developmental stage. The value for chromatin accessibility for each gene was defined as the sum of the average normalized peak counts of all the peaks associated with a gene at each studied region. The chromatin accessibility of the promoter region strongly correlated with the RNA expression at each stage (Figure 2C and 2D). Also, the changes in chromatin accessibility at the promoter region between E21 and P11 strongly correlated with the change in RNA expression between E21and P11 (Fig 2E). In contrast, the chromatin accessibility at the genic and distal region only weakly correlated to RNA expression (Supplementary Figure 3A-F). OCR visualization of selected RAGs, which were differentially expressed at the mRNA expression level between E21 and P11 (i.e. *Smad1, Myc, Jun, Klf2,* and *Klf9*), confirmed that the promoter region is more accessible in the stage with higher mRNA expression (Figure 2F and Supplementary Figure 4).

We next identified OCRs that are significantly more accessible at E21 than P11 or vice-versa. We identified, 15,402 differentially opened regions (DORs) at embryonic stage (E21 DORs), and 22,444 at postnatal stage (P11 DORs; Figure 2A). Interestingly, E21 DORs were enriched at promoters (<3 kb from TSS) when compared to P11, whereas P11 DORs were enriched in introns compared to E21 (Figure 3A, B and C). In the promoter region (<3 kb from TSS), there were 1,583 E21 DORs and 747 P11 DORs (Fig. 3D). Genes associated with these peaks were identified, resulting in 1,427 genes associated with E21 DORs (Fig 3E) including RAGs such as *Smad1, Tubb3, Jun, and Myc* (Figure 2F and Supplementary Figure 4). We found 689 genes that were associated with P11 DORs including RAGs such as *Klf9* and *Klf2* (Figure 2F and Supplementary Figure 4). Interestingly, only 29 genes were associated with both E21 and P11 DORs in the promoter region, with chromatin accessible sites changing in opposite directions during development (Fig 3E).

**Figure 3.**
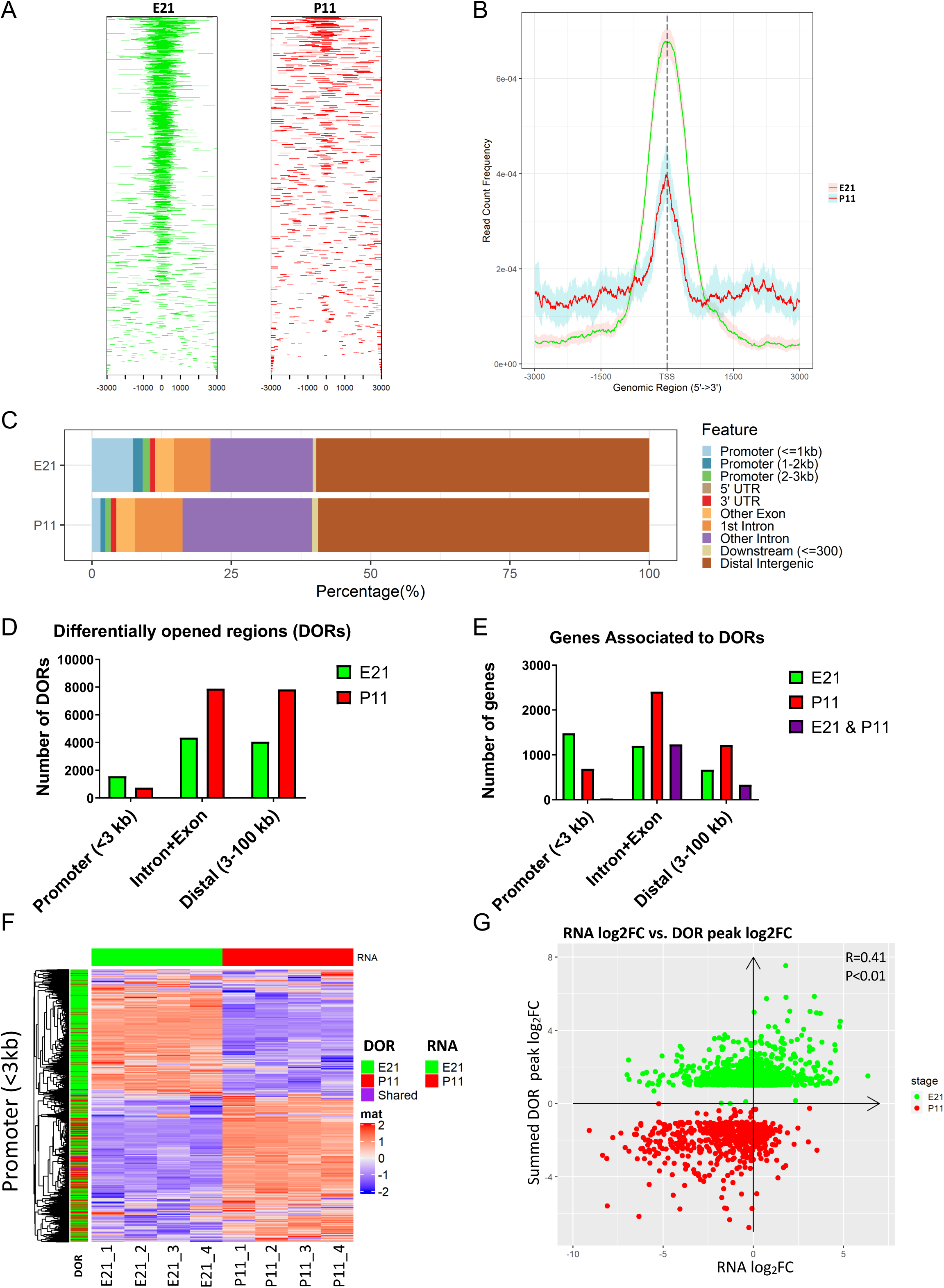
Differentially open regions are preferentially located at the promoter region of E21 genes and correlate with RNA gene expression changes. A) Distribution of DORs with respect the closest gene Transcription Starting Site (TSS). B) Read count frequency of DORs with respect to TSS. C) Distribution of DORs in the different regions of the genome. D) Number of DORs at the promoter region (<3 kb from TSS), distal region (>3 kb but <100 kb from TSS), and genic (intron+exon) region. E) Genes associated with DORs present at the promoter, distal, and genic region. F). RNA expression changes at E21 and P11 compared to the presence of DORs at the promoter region in E21 (green), P11 (red), or both (purple). G) Pearson correlation coefficient (p<0.01) of changes in gene expression between E21 and P11 and changes in DORs that are located in the promoter region.

To analyze the relationship of DORs with changes in RNA expression between E21 and P11, we plotted the RNA expression levels of each of the genes and the presence of stage-specific DORs (Figure 3F and Supplementary Figure 3G and I). We also performed a correlation analysis between the changes in RNA expression and the changes in chromatin accessibility of the DORs (Figure 3G and Supplementary Figure 3H and J)). We observed a strong correlation between changes of RNA expression and changes in chromatin accessibility in DORs at the promoter region (Figure 3F and G). In contrast, we observed a weak correlation at the genic region and no significant correlation at the distal region (Supplementary Figure 3G-J).

### Binding sites for TFs were identified in differentially opened regions of the chromatin and enriched at specific developmental stage

To identify potential TFs regulating the transcriptional change between E21 and P11, TF binding motifs were searched at DORs of either E21 or P11. We focused on chromatin region at promoters, since these regions strongly correlated with mRNA expression (Fig 3G). We calculated how much more probable it is to find a particular TF binding motif in stage specific DORs than in a random background set in the genome using the overrepresentation index (ORI), which takes into account the frequency and the density of a particular binding motif in a set of target sequences (Zhao *et al*, 2007) (Supplementary File 2). We identified six binding motifs that were significantly enriched at E21 DORs but not at P11 DORs (Fig 4A). These binding motifs include CREB/ATF, E74A (TF not present in mammals), bZIP911, v-Jun, ACAAT, and CCAAT. However, these last 4 binding motifs appear in less than 5% of the DORs undermining a widespread role regulating the E21 transcriptome. In contrast CREB/ATF appear in more than 50% of the DORs supporting an important role in regulating E21 genes (Supplementary File 2). We also found 78 binding motifs that appear significantly enriched in P11 DORs but not in E21 (Supplementary File 2). Thirty four of them with an ORI value higher than 1.5 appeared in more than 5% of DORs, such as MEF2, BACH, and RAR-related orphan receptors (Fig 4B). Binding motifs of certain TFs were found to be enriched in both E21 and P11 DORs suggesting that they control different genes depending on the cellular context. It is also noteworthy that some of these binding domains were more predominant in one particular stage as shown by the ORI ratio (Log2 E21 ORI / P11 ORI; Supplementary Figure 5 and Supplementary File 2). This might suggest that their activity or protein concentration is changing being higher in the developmental stage with higher ORI value. Some examples of these TFs include TAX/CREB, NRF-1, E2F, MYC, HIF1, and c-JUN in E21, and LXR in P11.

**Figure 4.**
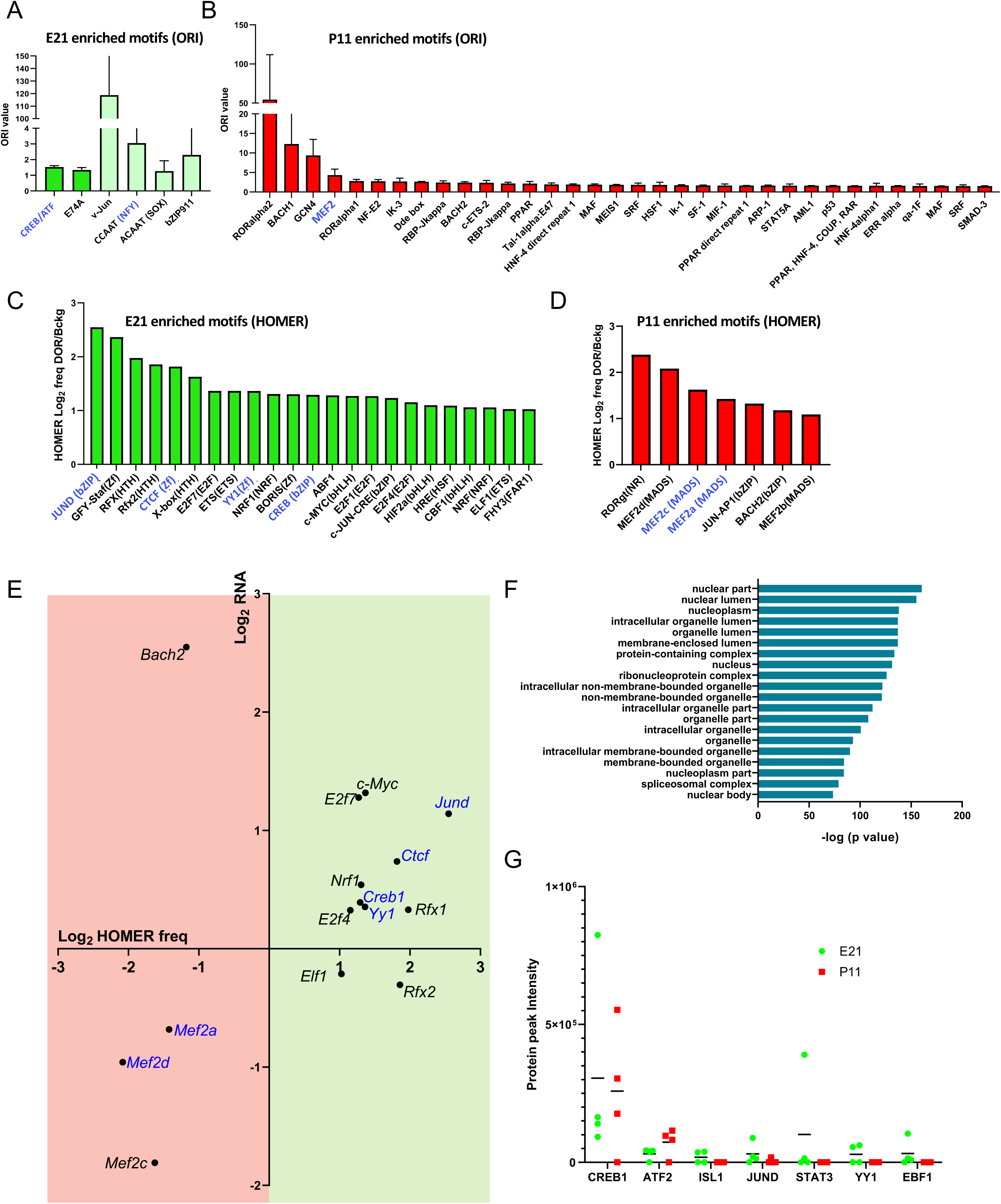
Binding site motifs of TFs expressed at the protein level are enriched at DORs of the promoter of RGC genes. A) Overrepresentation index (ORI) of TF binding motifs that are exclusively enriched at DORs of the promoter of E21 genes. Lighter green indicates binding motifs that appear in less than 5% of DORs. B) Overrepresentation index (ORI) of TF binding motifs that are exclusively enriched at DORs of the promoter of P11 genes. C) TF binding motifs that are exclusively enriched at DORs of the promoter of E21 genes according to the HOMER analysis. D) TF binding motifs that are exclusively enriched at DORs of the promoter of P11 genes according to the HOMER analysis. Only binding motifs that appear in more than 5% of all the DORs in either E21 or P11 and that appear in DORs with a frequency rate at least two times higher than the background set are plotted. E) TFs with binding motifs in C) and D) that appeared in RNAseq data were plotted over light green (E21) and light red (P11) background respectively. HOMER frequency ratio from C) and D) is represented together with the transcription factor RNA expression change between the two stages. HOMER frequency ratio in P11 was represented with a negative value. F) Cellular Component GO analysis of proteins identified in RGC nuclei by proteomics. G) Protein peak intensity of selected TFs (from A and B analysis) in E21 and P11 RGC nuclei isolated from rat independent litters (n=4 for each developmental stage). Protein peak intensity is normalized to total amount of protein of each sample. The black line represents the average of these samples in either E21 or P11 for each TF. Transcription factors in blue in A), B), C), D, and E) are those that were detected by proteomic analysis (LC-MS) in RGC nuclei. For overrepresentation index (ORI) of TF binding motifs, the p-value was calculated according to the method described in (Staden, 1989). Motif enrichment was calculated with cumulative binomial distributions statistics in HOMER.

We additionally performed binding site analysis using HOMER software (Supplementary File 3). In this case, the percentage of DORs containing the TF motif is calculated for E21 and P11 DORs and compared to a random background sequence to calculate statistical differences. Only TF motifs that appeared in more than 5% of DORs at the developmental stage with higher percentage, and that were twice as frequent compared to the background random set were shown (Fig 4C and D). We found binding motifs being enriched exclusively in E21 DORs such as CREB, c-MYC, CTCF, YY1, RFX, JUND, NRF1 and E2F (Fig 4C) and other set of binding motifs being enriched exclusively in P11 DORs, such as MEF-2, BACH2, and ROR receptors (Fig 4D). Other TFs that have been described as pro-regenerative in the literature such as SOX4, HIF1 and STAT3 were also significantly enriched in E21 DORs with frequencies 1.6 to 1.9 times higher than the background set (Supplementary File 3), which is lower than the cutoff we set in Figure 4C. Interestingly, CREB and MEF2 motifs were found to be uniquely enriched in E21 and P11 respectively, by both methods which utilize different TFBS motif database and very different statistical frameworks. Many TFBS motifs found to be more highly enriched in E21 than P11 by ORI ratio were found to be uniquely enriched in E21 by HOMER, e.g. MYC, JUND, NRF1 and E2F, demonstrating consistent high activity of those TFs in E21. The mRNA expression levels of most TFs correlated with their binding motif enrichment at their given developmental stage (Fig 4E) supporting the role of TFs, such as CREB, c-MYC, JUND, NRF1, CTCF, E2F7, RFX, YY1 and MEF2 in regulating RGC transcriptional change during development.

### Proteomic analysis of RGC nuclear fractions confirms the presence of TFs regulating transcriptional change

Since the presence of an mRNA does not always guarantee the presence of its corresponding protein, we performed a proteomic analysis of isolated nuclei from E21 and P11 RGCs to identify TFs that are expressed in RGCs at the protein level. A total of six E21 samples and five P11 samples were analyzed by LC-MS (Supplementary File 4). We identified 1192 proteins present in RGC nuclei. GO term analysis showed that these proteins were enriched in nuclear fractions (Fig 4F), confirming that nuclei isolation was efficient, in line with what was previously reported (Pita-Thomas *et al*., 2019). Uniprot analysis indicated that 72 of the proteins identified have DNA-binding activity and were detected at least in one sample. We identified several of the TFs that have enriched binding motifs in DORs such as CREB, JUND, c-MYC, CTCF, YY1, ATF2, MEF2a, and MEF2d (Fig.4A, B, C and D). We characterized the expression levels of these proteins by normalizing peak intensity for each TF in LC-MS to the total protein levels in the sample (Fig 4G). From all the TFs whose binding motifs are enriched at E21 DORs, CREB stood out for its consistent presence at high protein levels in RGC nuclei suggesting an important role in regulating transcription in RGC.

### CREB binding domain motifs are widespread at the promoter region of E21 DEGs

Since CREB and ATF2 are present at high protein levels in RGC nuclei and CREB/ATF binding motif (Fig 5A) is exclusively enriched at E21 DORs, we focused on this binding motif to identify the specific pathways regulated by CREB in E21 RGCs. From the 766 DEGs that have at least one OCR in their promoter, CREB/ATF binding motif appeared in 498 of them. We performed Biological Processes GO analysis for the group of genes with CREB/ATF binding motifs and found that these genes predominantly regulate neuron differentiation, axonogenesis and cytoplasmic translation (Fig 5B). We next performed Molecular Function GO analysis and found that CREB/ATF binding motif predominantly appears in the promoter of genes that have DNA binding transcription factor activity including regenerative TFs such as SMAD1, KLF7, MYC, or SOX11, and also ribosome genes (Fig 5C). GO Cellular Component analysis confirmed that many of the genes with CREB/ATF binding motif at their promoter region are ribosomal (Fig 5D). Altogether, these analyses suggest that CREB occupies a high rank in the hierarchical order of regenerative TFs and also plays an important role regulating ribosome biogenesis.

**Figure 5.**
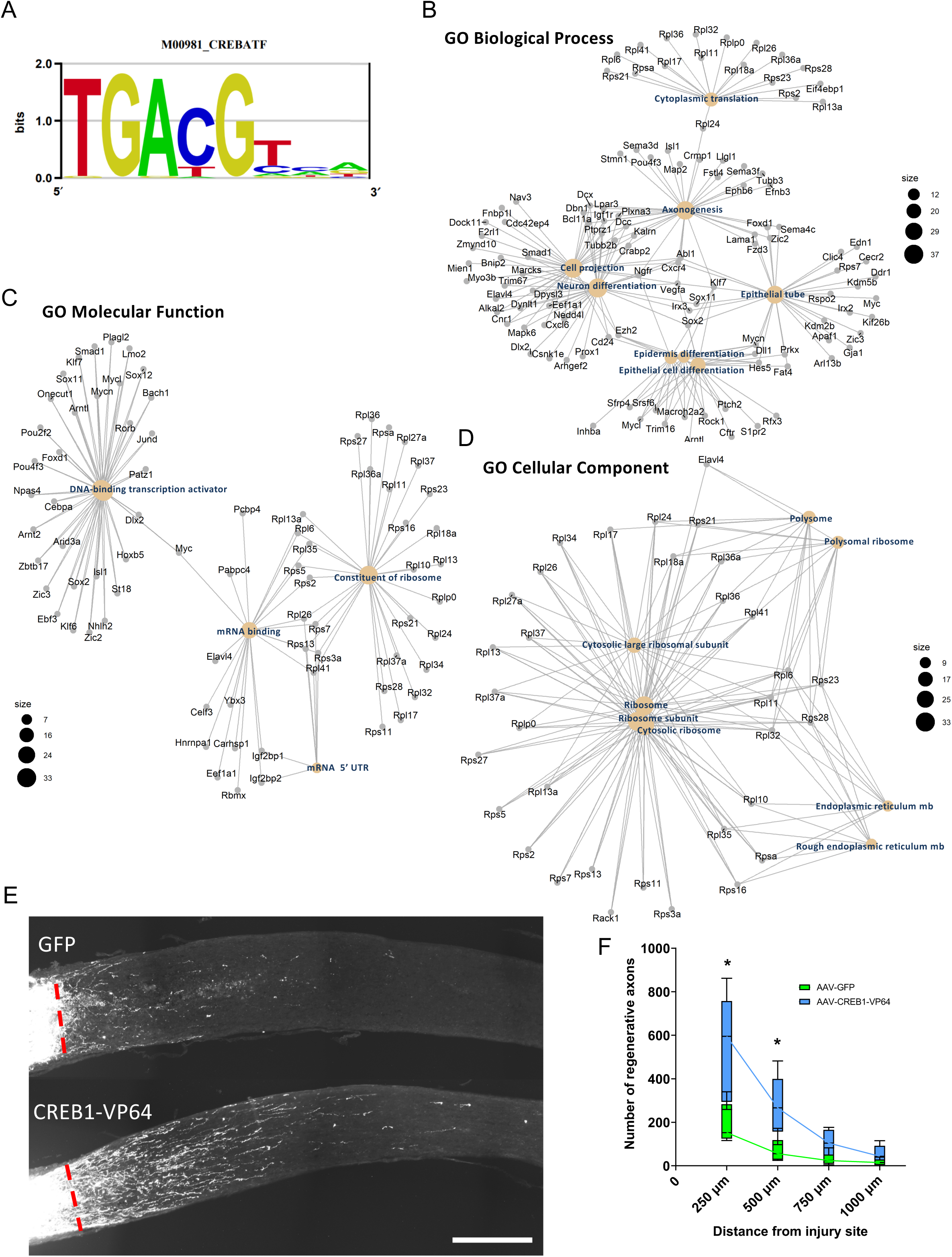
CREB1 regulates the changes in axon growth intrinsic capacity of RGCs during development. A) Sequence motif logo of the CREB/ATF position weight matrix. B) Gene hubs obtained by Gene Ontology analysis of E21 upregulated genes containing CREB/ATF binding motif at their promoter OCRs (For pathways analysis, FDR corrected P value < 0.05 were considered as significant.). Eight most significantly enriched Biological Processes are represented. C) Five most significantly enriched Molecular Functions are represented D) Eight most significantly enriched Molecular Functions are represented. E) Representative images of sliced optic nerves showing regenerative fibers of mice intravitreally injected with either AAV2-Creb1-VP64 or AAV2-GFP. Scale bar =250 μm. F) Number of regenerative fibers at various distances. Multiple t-test analysis with Holm-Sodak correction. * p adj <0.05. n=6 optic nerves per treatment. Four optic nerve sections were used to calculate the average for each nerve. Box-Whiskers plot was used where the box represents the 25th and 75th percentiles, the inside line represents the median, and the whiskers represents the largest and smallest values.

### CREB-VP64 overexpression promotes optic nerve regeneration

Our ORI and HOMER analysis revealed that CREB binding motifs are enriched in E21 chromatin accessible sites, suggesting that during development, the activity of this TF decreases as axon growth capacity declines. Since CREB mRNA and protein levels are highly expressed in RGCs but not drastically changing between E21 and P11, we tested if overexpressing an active form of CREB would promote axon regeneration. We fused CREB1 to the VP64 transactivation domain, which significantly increases the recruitment of transcriptional machinery (Beerli *et al*, 1998). VP64 is a tetramer of the 11 aa minimal activation domain of VP16, which results in increased activation compared to VP16 (Beerli *et al*., 1998). Previous studies have demonstrated that the addition of VP16 on pro-regenerative transcription factors significantly increases their ability to drive regeneration *in* vivo (Blackmore *et al*., 2012; Gao *et al*, 2004; Mehta *et al*, 2016). We performed intravitreal injections of AAV2-CREB-VP64 fusion, as described previously (Pita-Thomas *et al*., 2019). Fourteen days after viral injection, we performed optic nerve crush injury and allowed axons to regenerate for another fourteen days before injecting fluorescently labeled cholera toxin b to trace regenerating axons (Fig 5E,F). We found that overexpression of CREB1-VP64 significantly promoted optic nerve regeneration compared to GFP control, tripling the number of regenerating axons (Fig 5E,F). The extent of axon regeneration we observed two weeks after optic nerve injury is similar to what has been reported by manipulating SOX11, c-Myc or SOCS3 (Belin *et al*, 2015; Norsworthy *et al*., 2017; Smith *et al*, 2009) or by expressing the combination of the transcription factors Oct4-Sox2-Klf4 (Lu *et al*., 2020). These results indicate that CREB1-VP64 promote RGC axon regeneration, supporting a key role for the CREB1-dependent transcription program in the axon growth capacity of RGCs. Further studies will be needed to test if CREB1 overexpression synergizes with other strategies targeting the mTOR signaling pathways (Duan *et al*, 2015; Park *et al*., 2008; Pita-Thomas *et al*., 2019; Sun *et al*, 2011) or DNA methylation patterns (Lu *et al*., 2020).

## DISCUSSION

In the present study, we used RNAseq, ATAC-seq, and proteomics analysis to identify the key TFs regulating RGC axon growth capacity during development. We have shown that the RNA expression of genes that control axon growth and ribosome biogenesis are downregulated during RGC development, and that the mRNA changes correlate with chromatin accessibility at the promoter region. We found that binding motifs of several TFs that appear at the protein level such as CREB, JUND, CTCF and YY1 are enriched at these chromatin regions. The CREB/ATF motif is especially consistent and widespread, and it is enriched at the promoter regions of other regenerative TFs, suggesting a high rank of CREB in the hierarchical order of TFs regulating axon growth. In accordance with this role, overexpressing CREB1 fused to the VP64 transactivation domain in RGCs promotes optic nerve regeneration to a similar extent as targeting the mTOR pathway (Duan *et al*., 2015; Park *et al*., 2008; Pita-Thomas *et al*., 2019; Sun *et al*., 2011), DNA methylation (Lu *et al*., 2020) or expressing the transcription factors Sox11, c-Myc or SOCS3 (Belin *et al*., 2015; Norsworthy *et al*., 2017; Smith *et al*., 2009), partially reverting the poor regenerative capacity of adult RGCs.

Our analysis provides a map of the chromatin accessibility in acutely purified retinal ganglion cells at two different stages of development. Since RGCs represent less than 0.5% of all the cells in the retina, our study is a more specific analysis of the TFs regulating axon growth than studies analyzing whole CNS tissue (Dhara *et al*, 2019; Xie *et al*., 2020). Specifically, we found that the downregulation of genes in postnatal stages, such as axon growth genes and ribosomal genes, correlates with specific promoter regions becoming less accessible. In contrast, genes that regulate synapse function are upregulated during development and their promoter regions become more accessible. Our results are consistent with the observation that in the cortex, there is a developmental change in promoter accessibility that correlates with gene expression (Venkatesh *et al*, 2016). Venkatesh et al. study demonstrates that the target genes of the key RAGs STAT3 and JUN, but not SOX11 and KLF6, become less accessible in the adult stage (Venkatesh *et al*., 2016). This study was however limited by using a heterogenous population of cortical cells, whose proportions are dynamically changing during development. Another advantage of our study is that we complemented our TF binding motif analysis with proteomic analysis of RGC nuclei. This allowed us to confirm the presence at the protein level of several of the TFs whose binding motifs were enriched at DORs such as CREB1, JUND, CTCF, YY1, and ATF2. We also observed TFs whose target regions are becoming less accessible during RGC development including TFs with known roles in axon regeneration, such as CREB, JUN, SOX4, c-MYC, CTCF and HIF1 (Belin *et al*., 2015; Cho *et al*, 2015; Moore *et al*., 2009; Norsworthy *et al*., 2017; Watkins *et al*, 2013) (Gao *et al*., 2004; Palmisano *et al*, 2019), as well as other TFs whose roles in axon regeneration have not been thoroughly studied such as YY1, NRF1 and RFX. Interestingly, KLF’s motifs were not found to be predominantly enriched at any stage. KLF6 and KLF7 are upregulated in embryonic stage, and KLF2 and KLF9 in postnatal (Moore *et al*., 2009), and these TFs have opposing effects in axon growth. Since TFs of the same family bind to similar motifs, it is possible that we were not able to distinguish whether the binding motif present at the DOR is from an axon growth promoting KLF (KLF6 and KLF7) or axon growth inhibitory (KLF9 and KLF2). We also found that the promoter regions of regenerative RAGs such as SMAD1, MYC and TUBB3 were less accessible in P11 RGCs compared to E21. In Zebrafish, a species that can successfully regenerate axons in the optic nerve, chromatin accessibility does not significantly change between non injured RGCs and regenerating RGCs (Dhara *et al*., 2019). This suggests that in zebrafish, the chromatin remains fairly open at the target genes of regenerative TFs, whereas in mammals, the chromatin closes during development, limiting the access of regenerative TFs to activate the expression of RAGs after optic nerve injury. Altogether, these results supports the notion that in mammals, axon growth decline might be explained by the limited accessibility of TFs to certain regions of the chromatin.

Our chromatin analysis also showed that the DORs at the promoter of postnatal genes are enriched for certain TF binding motifs, including MEF2. Interestingly, MEF2 isoforms were upregulated in postnatal stage in our RNAseq data, and MEF2A and MEF2D were also identified at the protein level in postnatal RGC nuclei. This suggests that MEF2 might have an important role in regulating RGC maturation and potentially axon growth inhibition. Previous studies have shown that when HDAC5 is not phosphorylated, it is translocated to the nucleus where it binds to MEF2, repressing MEF2 target genes (Lu *et al*, 2000). In our previous study, we showed that a non-phosphorylated form of HDAC5 promotes optic nerve regeneration by activating the mTOR pathway(Pita-Thomas *et al*., 2019). Although we could not confirm the presence of HDAC5 in RGC nuclei, it is possible that some HDAC5 was present in the nucleus, repressing MEF2 target genes and contributing to its effects in promoting optic nerve regeneration.

It is noteworthy that the expression levels of CREB do not change dramatically between E21 and P11 according to our RNAseq and proteomics data. However, our chromatin accessibility analysis allowed us to identify CREB motifs in DORs suggesting that this TF is regulating transcriptional changes in embryonic stage. CREB activity is regulated by cAMP levels and previous studies have shown that the levels of cAMP drop after birth during CNS and PNS neuron development (Cai *et al*, 2001). In rat RGC, cAMP levels are three times higher in E18 compared to P5 (Cai *et al*., 2001). This suggests that CREB is activated and regulating axon growth genes in embryonic RGCs. After birth, cAMP levels drop and therefore transcription of axon growth genes by CREB ceases. Interestingly, activating CREB by increasing cAMP levels *in vivo* was not sufficient to promote optic nerve regeneration, but when combined with other stimuli such as oncomodulin or PTEN deletion, cAMP had a significant effect on optic nerve regeneration (Kurimoto *et al*, 2010). Similarly, elevating cAMP in cultured RGCs by adding forskolin does not revert the axon growth decline observed in postnatal RGC (Goldberg *et al*., 2002b). Here, we observed that the chromatin in the promoter region of axon growth promoting genes is less accessible in postnatal RGCs. In adult RGCs, endogenous CREB activated by cAMP may be unable to effectively access these chromatin sites to promote regeneration (Venkatesh *et al*., 2018). Transactivation domains such as VP16 and VP64 are well-known chromatin-modifying agents that open chromatin and greatly activate transcription (Tumbar *et al*, 1999). It has been previously shown that fusing VP16 to KLF7 is required to induce effective axon growth in adult RGCs by KLF7 (Blackmore *et al*., 2012). CREB-VP16 also promotes some level of axon regeneration in sensory neurons after spinal cord injury (Gao *et al*., 2004). However, whether CREB can promote regeneration in CNS neurons had not been tested. Here we demonstrate that expression of CREB-VP64 in adult RGCs promotes optic nerve regeneration. CREB-VP64, but not endogenous CREB activated by cAMP, might be able to open the chromatin of CREB target pro-regenerative genes inducing optic nerve regeneration.

Our study supports the role of CREB as a master regulator of axon regeneration. CREB binding motifs appear in the promoter DORs of RAGs that are downregulated during development. We found that the CREB/ATF motif is particularly enriched in signaling pathway that control axon growth, suggesting that besides CREB, other members of the CREB/ATF family such as CREM or ATFs could contribute to promote optic nerve regeneration. CREB/ATF also appear in DORs of ribosome genes which can also help to increase protein translation which has been shown to promote regeneration (Abe *et al*, 2010; Carlin *et al*, 2019; Park *et al*., 2008). More importantly, CREB binding motifs are enriched in OCRs at the promoter region of well-known regenerative TF genes such as SMAD1, MYC, KLF6, KLF7 and SOX11. This suggests that CREB occupies a high rank in the hierarchical order of regenerative TFs and that CREB sits atop several regenerative pathways. However, the extent of regeneration we observed with CREB-VP64 is similar to what has been shown with AAV-Sox11 and AAV-Myc (Belin *et al*., 2015; Norsworthy *et al*., 2017). It is possible that with a higher expression level of CREB, regeneration would have been stronger. For example, Cre inducible expression of c-MYC induces substantially more regeneration than when c-MYC is overexpressed using AAVs (Belin *et al*., 2015). Furthermore, although VP64 is an effective chromatin modifier (Tumbar *et al*., 1999), CREB may be unable to drive expression of certain target genes in closed chromatin. The activation of additional axon growth signaling pathways is likely required to reach CREB full transcriptional potential. For example, in cultured RGCs elevating cAMP with forskolin synergizes with the addition of growth factors such as CNTF or BDNF to the medium (Goldberg *et al*, 2002a). Finally, the presence of TFs that inhibit axon growth or the lack of other regenerative TF might limit the capacity of CREB to promote axon growth. Combining CREB-VP64 with other treatments targeting the neurotrophic factor CNTF (Muller *et al*, 2007; Pernet *et al*, 2013), mTOR (Duan *et al*., 2015; Park *et al*., 2008; Pita-Thomas *et al*., 2019; Sun *et al*., 2011) or DNA methylation (Lu *et al*., 2020) might provide a synergistic effect in optic nerve regeneration.

## MATERIAL AND METHODS

### Retinal Ganglion Cell Immunopanning Purification

For embryonic RGCs, E21 day pregnant Sprague Dawley rats were euthanized by CO_2_ and embryos were transferred to a plate with DPBS where retinas were dissected out and cleaned. For postnatal RGCs, P11 Sprague Dawley pups were euthanized by CO_2_ and eyes were transferred to DPBS medium where retinas were dissected. Next, RGCs were isolated as previously described (Pita-Thomas *et al*., 2019). Briefly, retinas were transferred to a filtered solution of papain in DPBS containing 2 mg of L-Cysteine and 2,000 units of DNAse. For E21 retina, 140 units of papain were added. For P11, 200 units were added. After 30 minutes, enzymatic solution was removed and retinas were triturated to obtain a single cell solution. RGCs were isolated by immunopanning. First by negative selection using an anti-macrophage antibody and then positive selection using anti-thy1 antibody. RGCs attached to the plate were trypsinized, counted in a Neubauer, and centrifuged for subsequent processing.

### RNA Sequencing and Analysis

Total RNA from 700,000 to 1.25 million RGCs from 4 independent rat litters per developmental stage was isolated using the Qiagen RNAeasy kit from Qiagen, which included a 15 minutes On-column DNAse step. RNA was stored at -80°C and sent to Genome Technology Access Center at Washington University for library preparation and sequencing. RNA quality was assessed using an Agilent Bioanalyzer (RIN > 9.5). Samples were subjected to DNase treatment. rRNA depletion was achieved with the Ribo-Zero rRNA removal kit. Library preparation was performed using the SMARTer kit (Clontech), and sequencing performed on an Illumina HiSeq3000. Basecalls and demultiplexing were performed with Illumina’s bcl2fastq software and a custom python demultiplexing program with a maximum of one mismatch in the indexing read. Sequences were adapter-trimmed using Cutadapt 1.16 (Martin, 2011) and subjected to quality control using PRINSEQ 0.20.4 (Schmieder & Edwards, 2011) and aligned to Rat (Rattus norvegicus) annotations based on genome assembly RNOR6 using STAR 2.5.3a (Dobin *et al*, 2013). Reads in features were counted using HTseq 0.6.1 (Anders *et al*, 2015).

Genes differentially expressed between conditions were identified using DESeq2 with log2FC > 1.0 and a false discovery rate (FDR) adjusted p values < 0.01, which includes a Benjamini-Hochberg correction (Love *et al*, 2014). Variance stabilizing transformation (VST) normalized counts were calculated using DESeq2, and normalized gene counts were converted to Z scores for plotting. Heatmaps were generated using ComplexHeatmap R package (Gu *et al*, 2016). Sequencing performance was assessed for total number of aligned reads, total number of uniquely aligned reads, genes and transcripts detected, ribosomal fraction, known junction saturation, and reads distribution over known gene models with RSeQC 2.6.24 (Wang *et al*, 2012). R package clusterProfiler (Yu *et al*, 2012) was used for GO and KEGG pathway enrichment analysis and plotting. GO and KEGG pathway terms with FDR corrected P value < 0.05 were considered as significant.

### ATAC-Seq

After isolation, 300,000 RGCs from two independent rat litters per developmental stage were centrifuged and resuspended in Neurobasal/B27+ medium and 10% DMSO, to quick frozen at -80 °C. Samples were processed by UCSD Center of Epigenomic Technologies using their proprietary assay for Transposase-Accessible Chromatin coupled with high-throughput sequencing (ATAC-seq). ATAC-seq Integrative Analysis Package (AIAP) was used for sequence quality assurance, mapping, peak calling, and downstream differential open chromatin analysis (Liu *et al*., 2020). Following data quality control analyses were performed for each sample and across the projects: (1) Reads under peak percentage ranged 31.6-39.3%. (2) Signal enrichments around the TSS relative to genome wide average, a metric which identifies datasets with high signal to noise ratios, ranged 10.8-14.4%. (3) Pearson correlation coefficient between the two replicates were calculated to measure the concordance between the two biological replicates.

### Transcription factor binding motif enrichment analysis and CREB target analysis

To identify candidate transcription factors that regulate differentially expressed genes, the HOMER (Heinz *et al*, 2010) and motif over-representation index (Motif-ORI) (Zhao *et al*., 2007) algorithms were used to identify transcription factor binding sites enriched in the differentially open region compared with random sampled sequences in the genome that are not in the open chromatin regions but has the same length distribution. The Patser program calculates the probability of observing a sequence with a particular score or greater (Hertz & Stormo, 1999; Staden, 1989) for the given matrix and determines the default cutoff score based on that P-value. Sequence of peaks were retrieved using BEDTools (Quinlan & Hall, 2010) and was scanned using Patser to identify CREB binding sites. Any gene with at least one CREB binding site in one of the gene-associated peaks was counted as target gene.

### RGC Nuclei proteome characterization by LC-MS

Nuclei from 700,000-1.25 million RGCs from 6 E21 and 5 P11 independent rat litters were isolated using the Thermo Scientific NE-PER Nuclear and Cytoplasmic Extraction Kit. Next, trypsin digestion was perform by adding a buffer containing 100mM ammonium bicarbonate, 10mM TCEP and 25mM iodoacetamide followed by digestion with trypsin at 37 °C overnight. The digested sample was acidified with 1%TFA then cleaned up with C18 tip. The extracted peptides were dried down. Proteomic samples were fractionated in a StageTip casted with four SDB-RPS disks (3M Empore SPE disks). Peptides were sequentially eluted with four buffers of increasing salt content. The four fractions were dried under vacuum and dissolved with 0.1 % formic acid for LC-MS analysis. Fractions of each sample were analyzed by LC-MS with a Dionex RSLCnano HPLC coupled to an Orbitrap Fusion Lumos (Thermo Scientific) mass spectrometer using a 2h gradient. Peptides were resolved using 75 µm x 50 cm PepMap C18 column (Thermo Scientific). Sequence mapping and label-free quantification were achieved using MaxQuant (version 1.6.1). MaxQuant was set up to search Human reference proteome (Uniprot.org). The digestion enzyme was set as trypsin. Carbamidomethylation of cysteine was set as fixed modification. Oxidation of methionine and acetylation of N-terminal of protein were specified as variable modifications.

### Optic Nerve Regeneration Assay

All surgical procedures were completed as approved by Washington University in St. Louis School of Medicine Institutional Animal Care and Use Committee’s regulations. Five week old C57Bl/6 mice were anesthetized with isoflurane and intravitreally injected with 1.25 µl of the AAV-CREB-VP64 or AAV-GFP virus two weeks before optic nerve crush using a Hamilton syringe and a pulled glass attached by an adaptor. Optic nerve crush was performed by a surgeon blinded to treatment as previously described (Pita-Thomas *et al*., 2019). Mice were anesthetized by isoflurane inhalation, and optic nerve was exposed and crushed for 3 seconds with a 55 forceps. For analgesia, 1mg/kg buprenorphine SR-LAB (ZooPharm) was administered subcutaneously. Two weeks after surgery, mice were intravitreally injected with 1.5µl of 1 mg/ml fluorescent Cholera toxin B. Two days after, animals were sacrificed by CO_2_ inhalation and perfused with PBS followed by 4% Paraformaldehyde. Eyes and nerves were dissected out and post-fixed for 4 hours, washed in PBS and incubated in 30% Sucrose solution overnight at 4 Celsius. Nerves were then dissected and sectioned in the cryostat at 11µm of thickness. Optic nerve sections were imaged in a florescent microscope. Number of axons growing at 250, 500, 750 and 1000 µm distal from the injury site were quantified by a researcher blinded to treatment. Multiple t test analysis with Holm-Sidak correction was performed.

## ACKNOWLEDGMENTS

We would like to thank members of the Cavalli lab for valuable discussions. This work was funded in part by a by NIH grants NS096034 and EY029077, and by the Stein Research to Prevent Blindness Innovation Award to V.C.

## AUTHOR CONTRIBUTIONS

W.P.T and V.C designed research; W.P.T. performed RGC purification and optic nerve regeneration assay. T.M.C analyzed RNA sequencing data. G.Z performed ATAC-seq and transcription factor binding site analyses. W.P.T and G.Z analyzed the data. W.P.T, G.Z and V.C wrote the manuscript. All authors approved the manuscript.

## CONFLICT OF INTEREST STATEMENT

The authors declare no competing interests

## DATA AVAILABILITY

The raw FASTQ files were deposited at the NCBI GEO database under the accession number GSE163564. The following secure token has been created to allow review of record GSE163564 while it remains in private status: yfupygcqhpwznif.

## SUPPLEMENTARY DATA

**Supplementary File 1.** RNA seq analysis

**Supplementary File 2.** Promoter 3kb ORI analysis

**Supplementary File 3.** Promoter 3kb Homer analysis

**Supplementary File 4.** Proteomics analysis

## SUPPLEMENTARY FIGURES

**Supplementary Figure 1.**
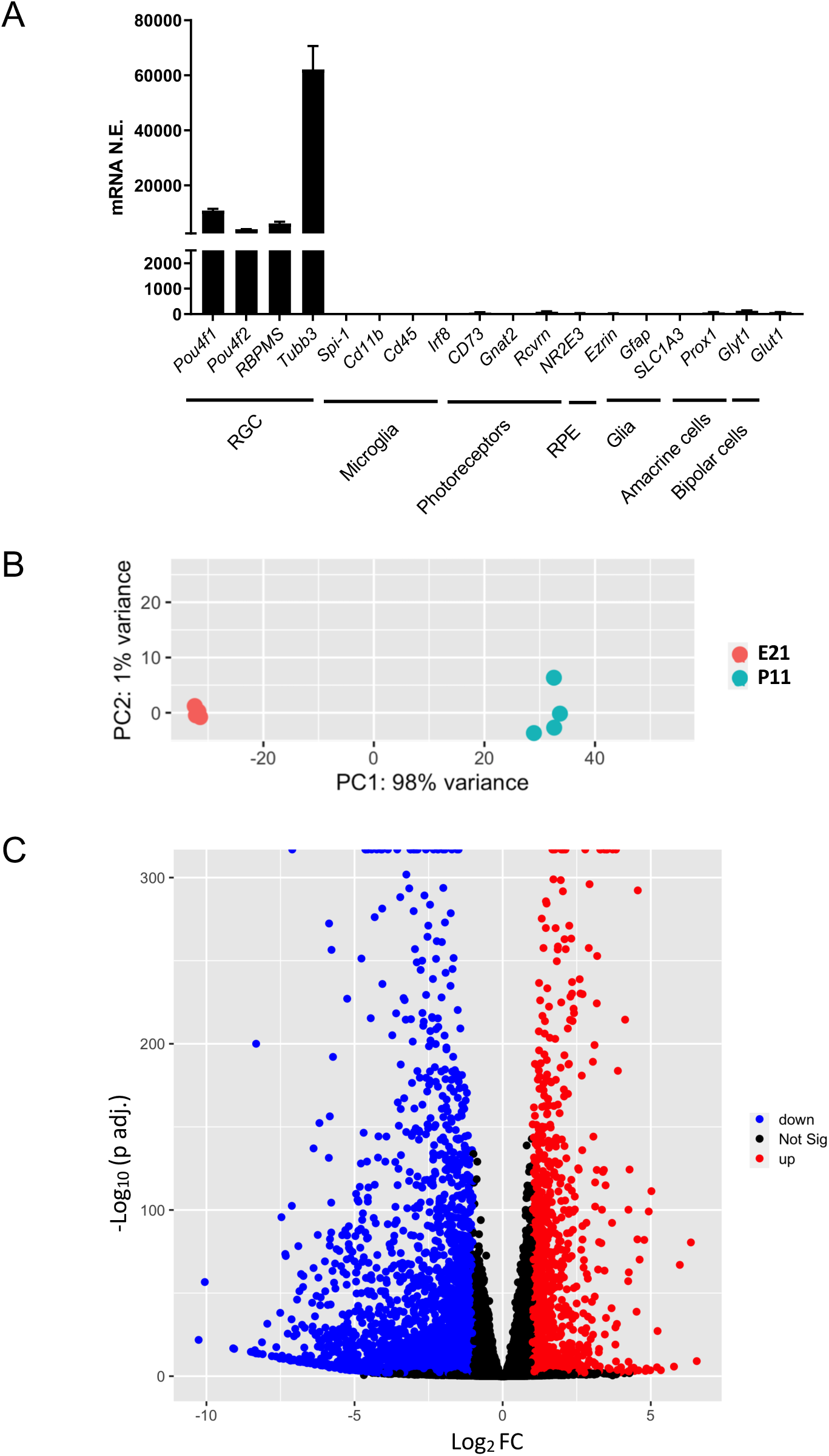
RNAseq data of E21 and P11 RGCs. A) The average (± SEM) RNA expression of RGC and other retinal cell markers in the 8 RGC samples from independent rat litters (E21 + P11). B). PCA analysis of E21 and P11 RNAseq samples. C) Volcano plot representing fold change between E21 and P11 and p-value. Cut off is Log2 FC>1 and false discovery rate (FDR) adjusted p<0.01. Blue, genes downregulated in P11, and, red, genes upregulated in P11.

**Supplementary Figure 2.**
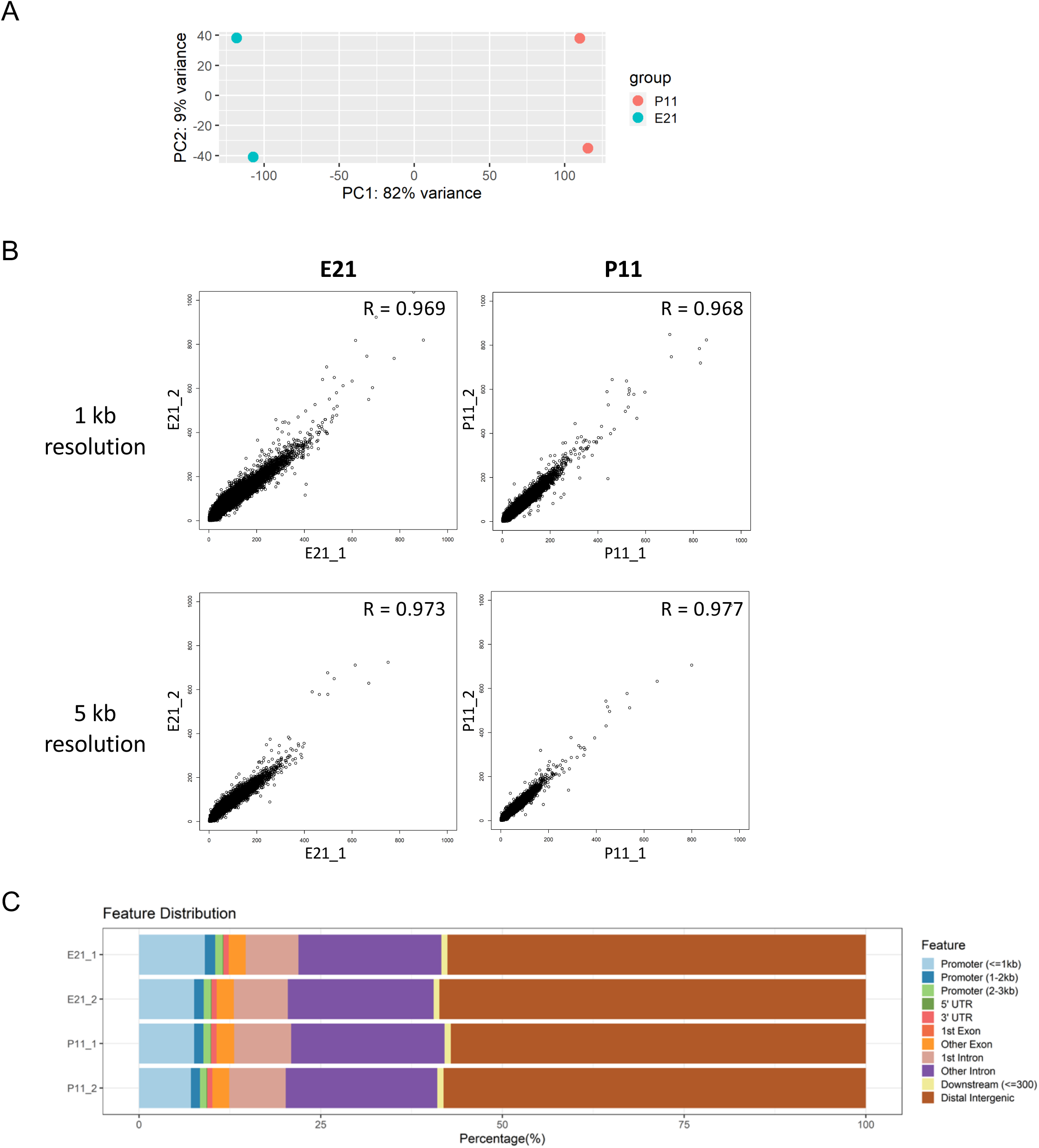
ATAC-seq analysis of E21 and P11 RGCs. A) PCA analysis of chromatin accessible sites of E21 and P11 samples. B) Pearson correlation coefficient analysis of chromatin accessibility signal in the same stage samples at 1kb and 5 kb resolutions. C) Proportion of chromatin accessible sites in the different genome locations in E21 and P11 RGCs.

**Supplementary Figure 3.**
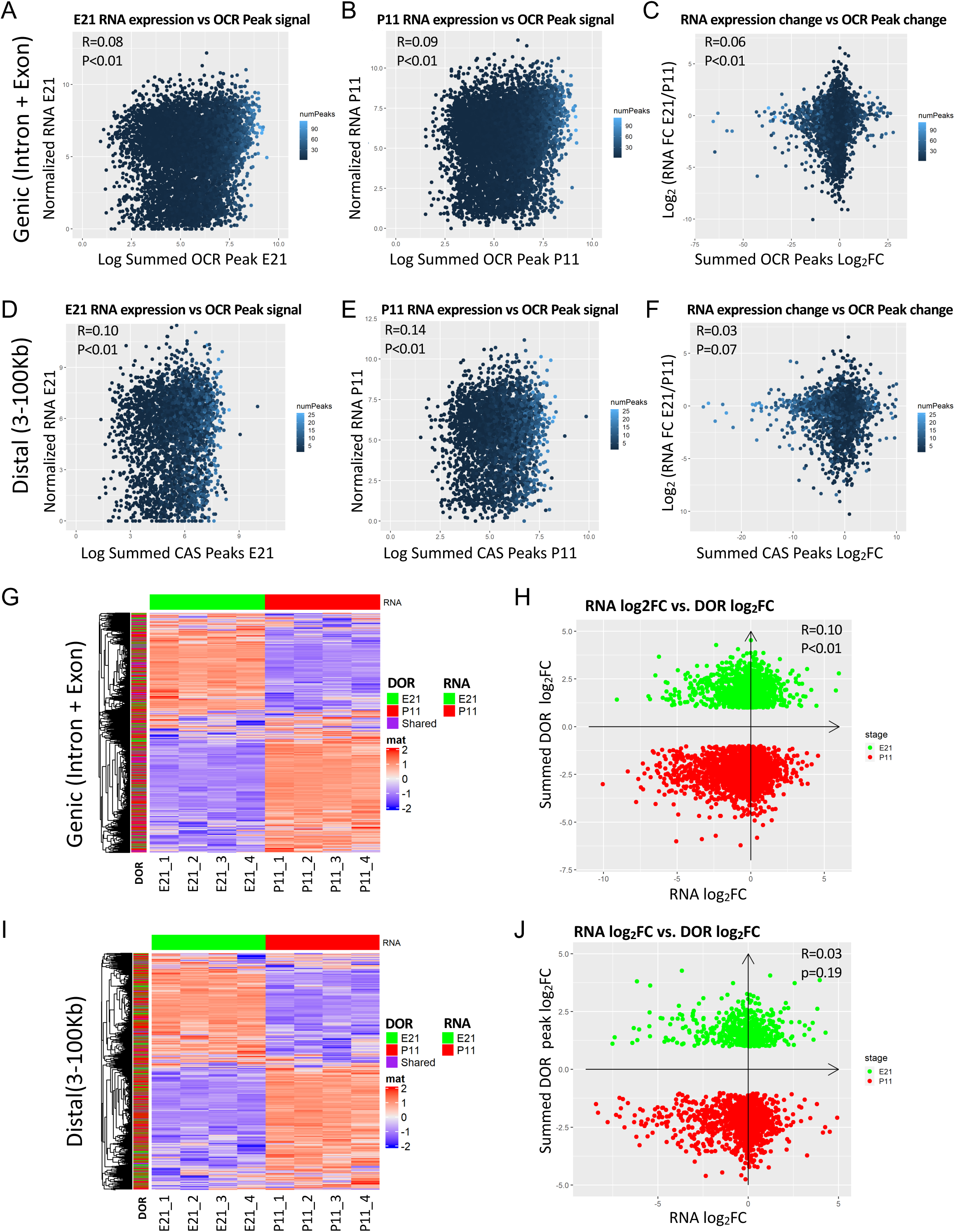
Correlation between RNA expression and chromatin accessibility in RGCs. A) Pearson correlation coefficient (p<0.01) between E21 RNA expression and chromatin accessibility (sum of all peak values) at the genic region (intron + exon) of genes in E21. B) Pearson correlation coefficient (p<0.01) between P11 gene RNA expression and chromatin accessibility at the genic region (intron + exon) of genes in P11. C) Pearson correlation coefficient (p<0.01) between the change in RNA expression between E21 and P11 and the change in chromatin accessibility at the genic region (intron + exon) of genes between those stages. D) Pearson correlation coefficient (p<0.01) between E21 RNA expression and chromatin accessibility (sum of all peak values) at the distal region (>3 kb but <100Kb) of genes in E21. E) Pearson correlation coefficient (p<0.01) between P11 gene RNA expression and chromatin accessibility at the distal region of genes in P11. F) Pearson correlation coefficient (p<0.01) between the change in RNA expression between E21 and P11 and the change in chromatin accessibility at the distal region of genes between those stages. G). RNA expression changes at E21 and p11 compared to the presence of DORs at the genic region in E21 (green), P11 (red), or both (purple). H) Pearson correlation coefficient (p<0.01) of changes in gene expression between E21 and P11 and changes in DORs that are located in the genic region. I). RNA expression changes at E21 and p11 compared to the presence of DORs at the distal region in E21 (green), P11 (red), or both (purple). J) Pearson correlation coefficient (p<0.01) of changes in gene expression between E21 and P11 and changes in DORs that are located in the distal region.

**Supplementary Figure 4.**
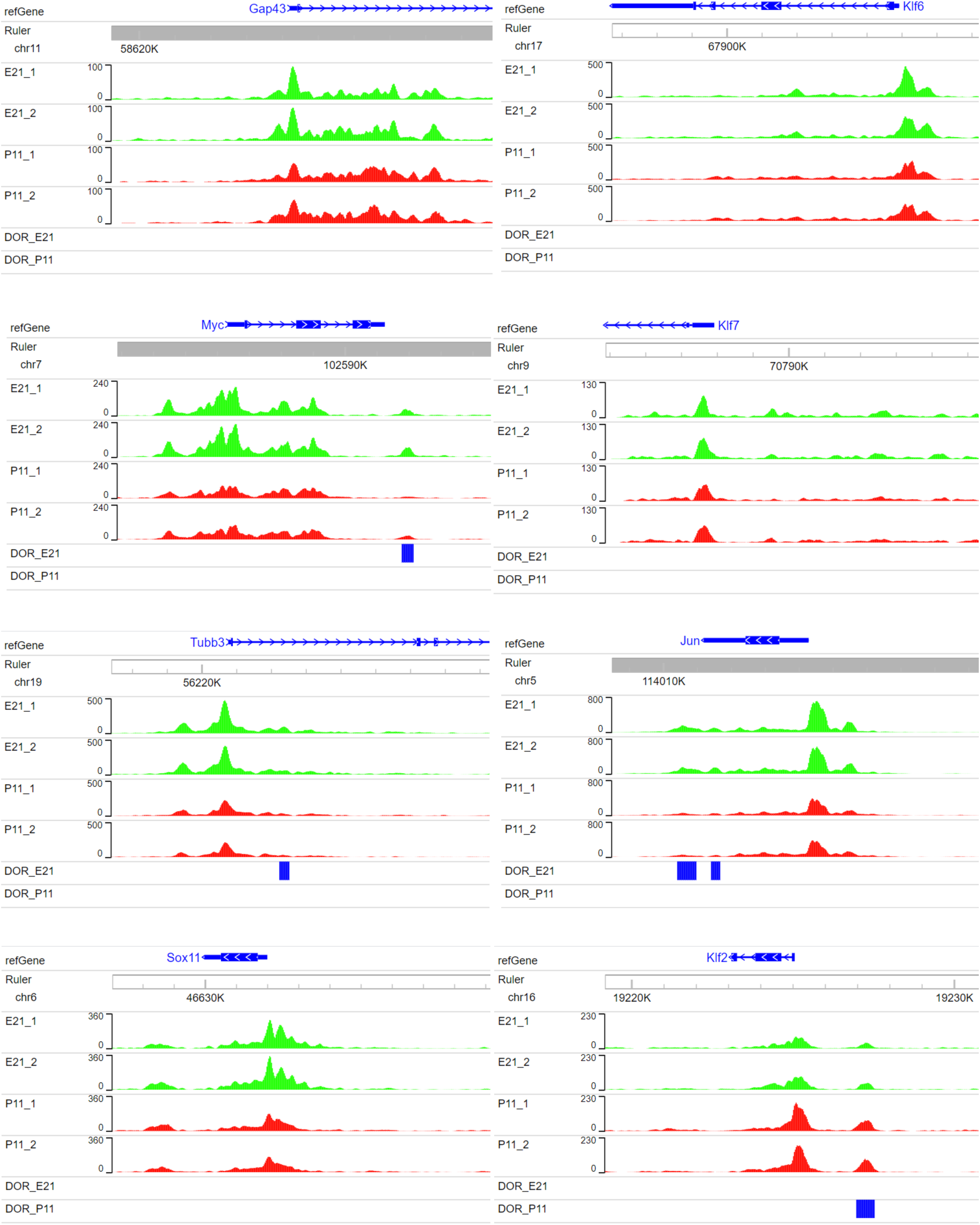
Visual representation of chromatin accessibility of E21 and P11 replicates near the TSS of regenerative associated genes *GAP43, Myc, Tubb3, Jun, Klf6, Klf7,* and *Sox11* (promoters of axon growth) and *Klf2* (inhibitor of axon growth). DORs are represented in blue accordingly to the developmental stage where this region is more accessible.

**Supplementary Figure 5.**
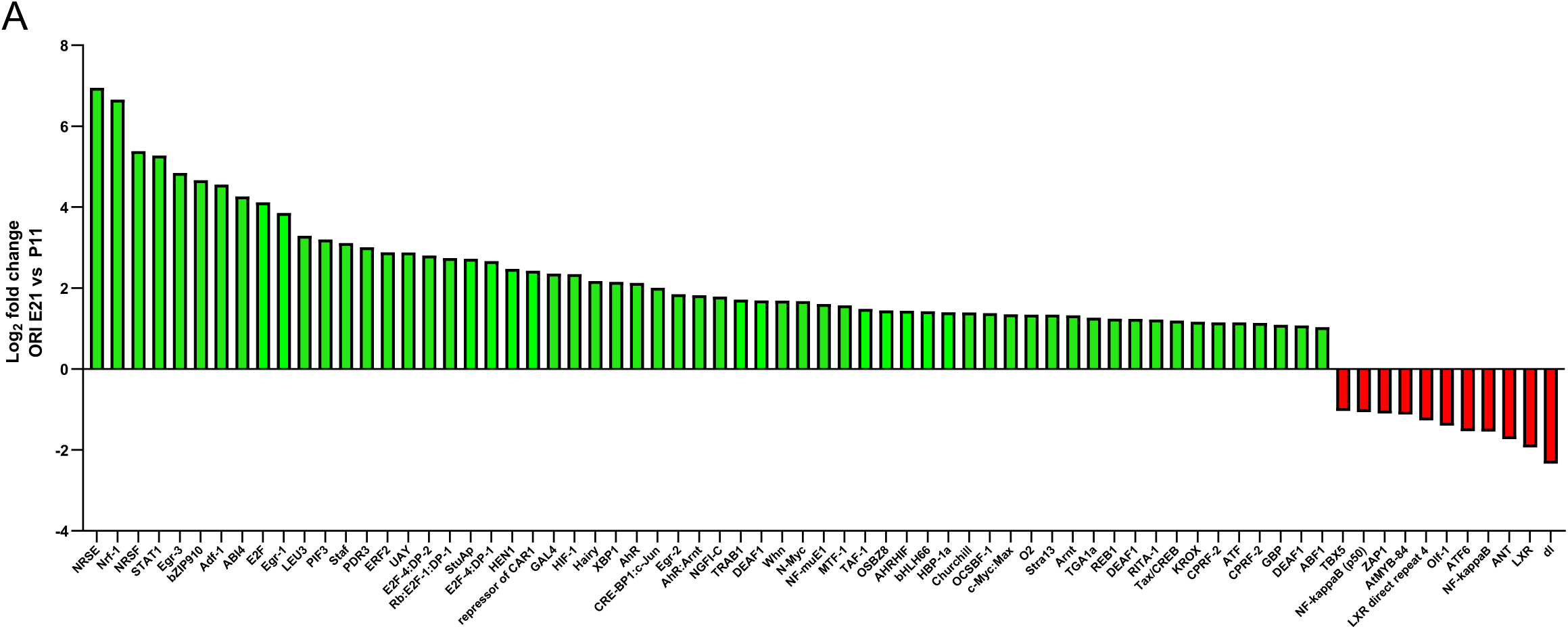
Binding Motifs of TFs that appear at both E21 and P11 DORs of the promoter. A) TF binding motifs with an overrepresentation index (ORI) significantly different to a background random set in both E21 and P11 DORs of the promoter. The ratio of the E21 ORI value/P11 ORI value represents the enrichment of a TF binding motif in a particular stage compared to the other developmental stage. Only TF binding motifs with a log_2_ (E21 ORI/P11 ORI)>1 or <-1 are plotted. Some TFs have multiple position weight matrices enriched and the one with highest absolute ORI ratio value was plotted.

